# Reconstruction of the molecular evolution of Usutu virus in Germany: Insights into virus emersion and circulation

**DOI:** 10.1101/2023.02.28.530558

**Authors:** Felicitas Bergmann, Cora M. Holicki, Friederike Michel, Sabine Bock, Nelly Scuda, Grit Priemer, Susanne Kenklies, Timo Siempelkamp, Jasmin Skuballa, Claudia Sauerwald, Louise Herms, Aemero Muluneh, Martin Peters, Andreas Hlinak, Martin H. Groschup, Balal Sadeghi, Ute Ziegler

## Abstract

Usutu virus (USUV) is a mosquito-borne flavivirus that is widely distributed in southern and central Europe. The zoonotic virus circulates primarily between birds and mosquitoes, can, however, in rare cases infect other mammals including humans. In the past USUV has been associated with mass mortalities in birds, formerly blackbirds and owls. Birds commonly succumb either due to the peracute nature of the infection or due to severe encephalitis. In Germany, USUV has spread rapidly since its first detection in 2010 in mosquitoes under the presence of susceptible host and vector species.

Nonetheless, there is to date limited access to whole genome sequences resulting in the absence of in-depth phylogenetic and phylodynamic analyses. In this study, 118 wild and captive birds were screened using a nanopore sequencing platform with prior target enrichment via amplicons. Due to the high abundancy of Europe 3 and Africa 3 in Germany an ample quantity of associated whole genome sequences was generated and the most recent common ancestor could be determined for each lineage. The corresponding clock phylogeny revealed an introduction of USUV Europe 3 and Africa 3 into Germany three years prior to their first isolation in the avifauna in 2011 and 2014, respectively. Based on the clustering and temporal history of the lineages, evidence exists for the genetic evolution of USUV within Germany as well as new introductions thereof into the country.

## Introduction

Usutu virus (USUV) is an arbovirus which belongs to the *Flaviviridae* family, genus *Flavivirus*. Its positive sense single stranded RNA genome, of 11,064 nucleotides, encodes a single polyprotein which is cleaved by viral and host proteases into three structural (C, prM, E) and seven non-structural proteins (NS1, NS2a, NS2b, NS3, NS4a, NS4b, and NS5) (1, 2). USUV was first isolated in Swaziland, Africa in 1959 from a mosquito (3). USUV circulates in an enzootic cycle between mosquitoes as vectors and birds as reservoir and amplifying hosts. Mosquitoes belonging to the *Culex pipiens* complex represent the main vector (4).

Extremely susceptible bird species such as passerine species including blackbirds (*Turdus merula*), or house sparrows (*Passer domesticus*), and birds of prey like great grey owls (*Strix nebulosa*) serve as natural hosts (5–8). So far, USUV was found primarily in association with die-offs of susceptible birds, formerly blackbirds in Germany (9). These mass mortality events have occurred in wild birds all over Central Europe (e.g., in Austria (7), Hungary (6), Switzerland (10), Italy (11), Czech Republic (12), and Germany (8)) since the first reported outbreak in Vienna, Austria in 2001 (13) (Fig 1). In retrospect, the first known occurrence in Europe dates back to 1996 in Italy, in association with a blackbird mortality event (14). However, Engel et al. 2016 (15) assumed that the virus was probably already introduced earlier, between the 1950s and 1960s, via bird migration from Africa into Western and Central Europe, followed by a rapid geographic spread of the virus.

**Fig 1.**
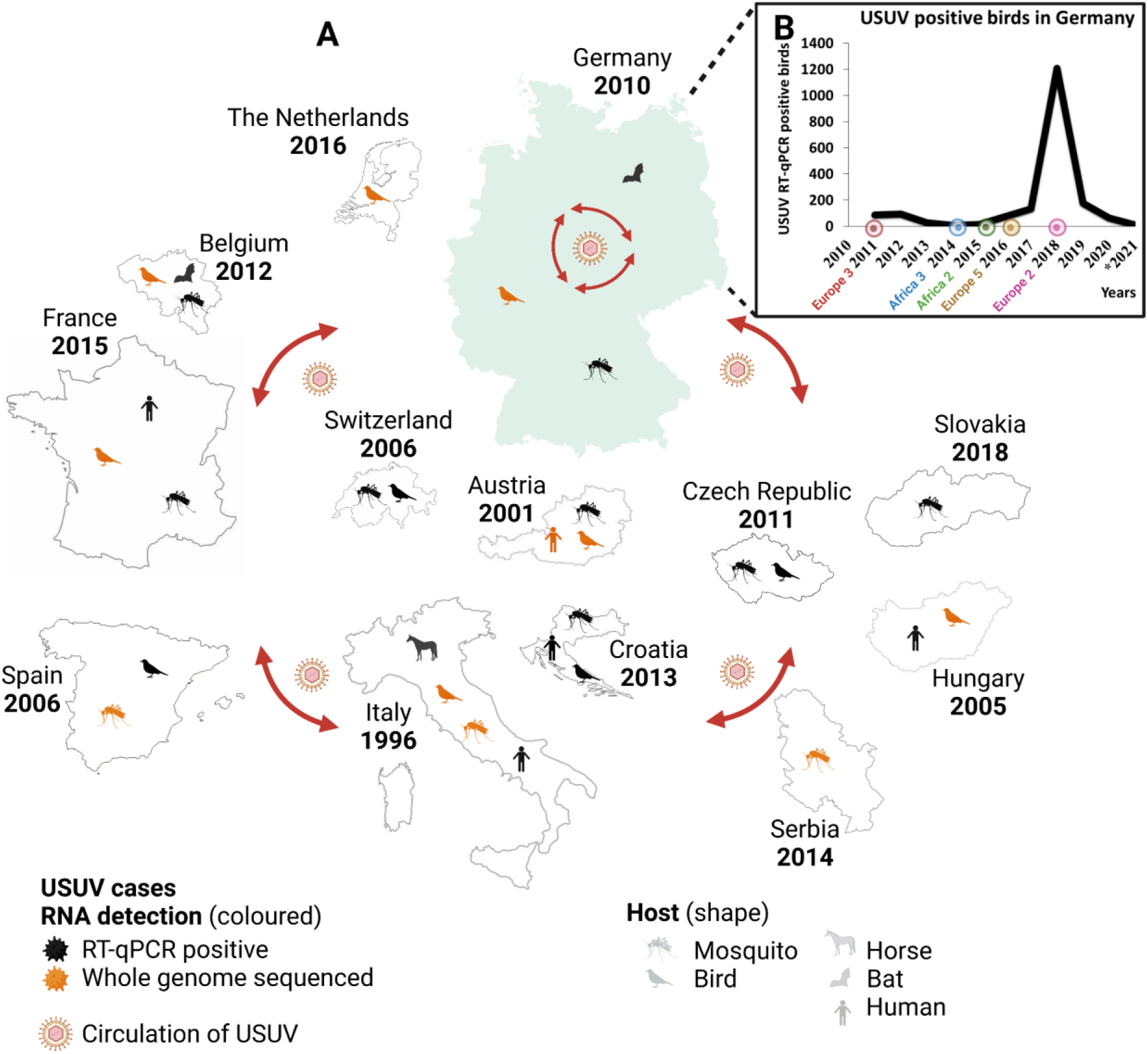
A) Geographic distribution of USUV-RNA detection throughout Europe. The first detected USUV-positive case per country is depicted in the figure. In addition, the icons for mosquito, bird, horse, bat, and human indicate in which species USUV-RNA has been detected so far and whether a corresponding whole genome sequence is available (in orange). (B) The graph shows all USUV-positive birds detected in Germany since 2011, highlighting the first occurrence of each lineage in the following years, with the exception of USUV Europe 3, which was first detected in 2010 in a mosquito pool. *Results not finalized and only based on dead bird surveillance (updated on 21^th^ February 2023). Created with BioRender.com.

The virus also has a zoonotic potential and can infect humans, in rare cases causing severe neurological symptoms (16–23). Humans as well as other susceptible mammalian species are considered dead-end hosts as they can become infected but cannot sustain the transmission cycle. USUV has in the past been found in horses (24, 25), dogs (26), deer (27), wild boar (28), bats (29), squirrels (30), and rodents (31). The genetic variability of USUV was studied by phylogenetic studies targeting partial sequences, especially of the envelope E and NS5 genes, and whole genome sequences (15, 32–35, 4, 36). These analyses resulted in the declaration of eight distinct lineages, which cluster together according to their geographic origin of detection: Africa 1-3 and Europe 1-5. Migratory birds from Africa are thought to have brought different USUV lineages to Europe at independent time points, resulting in the dispersal of distinct USUV lineages. As evidenced by phylogenetic analyses these lineages amplified and evolved independently. The genetic heterogeneity of the European lineages is, therefore, most likely due to in situ evolution rather than new introductions by long-distance migratory birds (15). By comparison, widespread migration patterns and multiple introductions of virus variants from different geographic areas of origin resulted in the African lineages (15). Nonetheless, in the literature the assignment and nomenclature of USUV lineages/strains has not been standardized. It has also not been conclusively determined whether the different lineages have an influence on host and vector affinity (15).

Circulating in Europe, USUV did not stop at the boundary to Germany (Fig 1 (A)). In Germany, initial surveillance efforts for flaviviruses focused on the serological detection of WNV antibodies (37, 38). USUV-antibodies were solely determined in the further clarification of flavivirus cross-reactivity (37). However, since 2009 molecular surveillance programs in mosquitoes and birds were implemented, followed by several introductions into the country (39–46). The first USUV isolate (lineage Europe 3) was documented in Germany in 2010 from a pool of *Culex pipiens* biotype *pipiens* mosquitoes in the south of Frankfurt, in Weinheim (39). Subsequently, fatal cases in wild and captive birds, mainly Eurasian blackbirds and owls, were reported from the north of the Upper Rhine valley and adjacent areas of the Palatinate and the Neckar valley to the Southwest of Germany (8, 43). Around the same time (2014), a new USUV lineage (Africa 3) was introduced into the north of Germany, in Bonn, where only one case in a blackbird was detected (47, 48). In Berlin in 2015, the lineage Africa 2 occurred for the first time in two juvenile great grey owls (47). In 2016, USUV (lineage Europe 3, Africa 3, and Africa 2) continued to spread, with numerous cases reported in the southwest, northwest, and east of Germany concurrent with the first detection of Europe 5 in central-western North Rhine-Westphalia (44, 49, 33). In addition, a further spread to neighbouring countries to the west could be confirmed (33). In the Federal State of Saxony in 2018, USUV lineage Europe 2 was detected for the first time (45). Until 2018, the virus had spread nationwide with five USUV lineages present (Africa 2, Africa 3, Europe 2, Europe 3, and Europe 5) (Fig 1 (B)).

To date, the circulation of USUV has been reported (based on serological and molecular evidence) in many countries in and around Europe: Tunisia (50), Morocco (51), Israel (52), Greece (53), France (54), Spain (55), Poland (56), Hungary (6), Czech Republic (57), Serbia (35), the United Kingdom (58), Croatia (59), the Netherlands (60), Switzerland (10), Italy (11), and Germany (39). Ongoing efforts to elucidate the USUV phylogenetic scenario in Europe, reported the co-circulation of Europe 3 and Africa 3 in the Netherlands (36), Europe 3 and Africa 2 and 3 in France (54), and Europe 1-3 and Africa 3 in the Czech Republic (61). In Germany, there is an evident co-circulation of the USUV lineages Europe 2 and 3, and Africa 2 and 3 in 2017 and 2018 (45). A recently published study based on partial sequences from 2019 and 2020, could confirm the ongoing circulation of USUV lineages Europe 2 and 3 as well Africa 3 in the country (46).

So far, only a small number of USUV whole genome sequences from Germany are publicly available, making it difficult to determine the precise time point of USUV introduction into the country. In addition, it is not clear whether the virus was introduced once or whether several independent introductions took place. Third-generation sequencing technologies like Nanopore MinION sequencing, a sequencing platform validated in this study, enable new and more accessible ways of studying infectious diseases. It can be used to clarify the origin, transmission routes, and ecology of emerging viral diseases, to tackle unanswered fundamental questions (62–66). Therefore, in this study 118 USUV genomes from wild and captive birds were sequenced using Nanopore MinION to further unravel the occurrence and spread of USUV in Germany from 2017 to 2021. Whole genome sequencing (WGS) produces a greater and more complete data set as compared to partial sequencing and is therefore a promising tool in gaining an in-depth understanding of USUV introduction events into Germany and in characterizing the evolutionary history of USUV.

## Material and Methods

### Samples

Blood and organ samples of 118 wild and captive birds were collected in the frame of ongoing monitoring bird studies and in close collaboration with the local state laboratories in Germany and submitted to the national reference laboratory (NRL) for West Nile virus (WNV) and USUV at the Friedrich-Loeffler-Institut (FLI). From 2019 to 2021, a total of 3,762 birds were tested for USUV. The samples were recorded in a database for the detection of USUV-RNA in birds, which was established in 2019 with the aim of providing the public health authorities with a nationwide overview (46). Unfortunately, prior to 2019, there was no comparable database and the number of tested birds in addition to those that were part of the study described by Michel et al. 2019 (45) can only be estimated for 2017 and 2018 and are most likely higher in reality (Table 1).

For WGS, USUV RNA-positive samples were selected primarily on the basis of their geographic location in order to represent a comprehensive picture of Germany. The cycle threshold (Ct) values and to some extent also the lineages, as already determined by partial sequencing (46, 45, 67), were used as further decision criteria. Throughout all five years, USUV isolates from the two different bird orders *Passeriformes* and *Strigiformes* were identified and sequenced. In 2019, USUV was additionally found and sequenced in *Anseriformes* and *Columbiformes* (Table 2). Viral RNA of blood and organ samples was isolated using the RNeasy^®^ Mini Kit (Qiagen, Hilden, Germany) according to the manufacturer’s protocol. Cell culture supernatant samples were extracted using the Viral RNA Mini Kit (Qiagen) following the manufacturer’s instructions. Analysis of extracted RNA was performed using real-time reverse transcription PCR (RT-qPCR) assays specific for USUV, as described by Jöst et al. 2011 (39). Samples with Ct values between 12.09 and 32.79, covering almost all Federal States of Germany, were included in this study. Detailed information of each sample is provided in S1 Table. Four of the sequenced birds (GenBank accession numbers: OP422562-OP422565) were already published in a next-generation sequencing (NGS) methodological publication by Holicki et al. 2022 (67).

**Table 1.** Number of molecularly tested birds per year, from the live and dead bird surveillance, including USUV RT-qPCR positive results and number of samples sequenced (68, 69, 45, 70).

| Year | Total Tested Birds | USUV RT-qPCR Positive | Samples Sequenced |
| --- | --- | --- | --- |
| 2017 | NA | 132 | 6 |
| 2018 | NA | 1,208 | 45 |
| 2019 | 2,202 | 177 | 37 |
| 2020 | 3,086 | 64 | 25 |
| 2021* | 1,383 | 12 | 5 |
| **Total** | **6,671** | **1,593** | **118** |

NA= not available
\*Results not finalized and only based on dead bird surveillance (updated on 21th February 2023).

**Table 2.** Overview of sample set.

|  | 2017 | 2018 | 2019 | 2020 | 2021 |
| --- | --- | --- | --- | --- | --- |
| **Total (Wild/Captive Birds)** | 6<br>(3/3) | 45<br>(34/11) | 37<br>(29/8) | 25<br>(22/3) | 5<br>(1/4) |
| **Species Order** | *Passeriformes*,<br>*Strigiformes* | *Passeriformes*,<br>*Strigiformes* | *Passeriformes*,<br>*Strigiformes*,<br>*Columbiformes*,<br>*Anseriformes* | *Passeriformes*,<br>*Strigiformes* | *Passeriformes*,<br>*Strigiformes* |
| **Organs** | Brain, CNS, liver, spleen | Blood, brain, CNS, liver, pool of organs, spleen | Blood, brain*, CNS, kidney, liver, pool of organs, spleen | Blood, brain*, pool of organs, RNA, spleen | Blood, brain, liver |
| **Federal States** | NW, NI, SN | BE, BY, MV, NI, NW, SL, ST | BB, BE, BY, HE, NI, NW, SL, SN, ST, TH | BB, BE, BW, HE, NI, SN, ST | MV, NI, ST |
| **Lineages** | Africa 3, Europe 3 | Africa 3, Europe 3 | Africa 3, Europe 3, Europe 2 | Africa 3, Europe 3, Europe 2 | Africa 3, Europe 3 |

\* Selected samples were either diluted 1:5 in phosphate buffered saline (PBS) or were passaged in cell culture prior to RNA extraction. Abbreviation: CNS: central nervous system; BB: Berlin Brandenburg; BE: Berlin; BW: Baden-Württemberg; BY: Bavaria; HE: Hesse; NI: Lower Saxony; MV: Mecklenburg-West Pomerania; NW: North Rhine-Westphalia; SL: Saarland; SN: Saxony; ST: Saxony-Anhalt; TH: Thuringia

### Nanopore sequencing and data analysis

Nanopore sequencing was performed as described by Holicki et al. 2022 (67). In short, RNA was reverse transcribed using the SuperScript IV First-Strand cDNA Synthesis Reaction Kit (Cat. no. 18091050; Invitrogen by Thermo Fisher Scientific, Darmstadt, Germany) with random primers as previously described by Quick et al. 2017 (71). Followed by an USUV-specific multiplex PCR which was performed with two separate mixes of primer pairs using AccuPrime Taq DNA Polymerase High Fidelity (Cat. no. 12346-086; Invitrogen). MinION sequencing was carried out following the manufacturer’s instructions using the 1D Native barcoding genomic DNA Kit (Nanopore, EXP-NBD104 and SQK-LSK109, Oxford Nanopore-technology (ONT)) on a Spot-ON flow cell (R9.4.1; ONT). Twelve samples were multiplexed per flow cell. Fast5 raw data reads were demultiplexed using Guppy v4.5.4 (72). Primers were trimmed and reads were quality controlled to a minimal length of 200 base pairs (bp) and reads with a minimum quality of 7 were considered for further analysis. For consensus sequence generation, an alignment against the selected USUV reference genomes v23 (73) was performed using KMA (74)and Minimap2 (75). Consensus sequences were visualized with Geneious Prime 2021.0.1 (Biomatters Ltd., Auckland, New Zealand).

### USUV genome phylogenetic analyses

USUV sequences of the National Center for Biotechnology Information (NCBI; Bethesda, MD, USA) were screened and all available full length USUV genomes from Germany up to August 24, 2022 were downloaded (76). All full length USUV genomes including the newly sequenced USUV genomes were aligned using MUSCLE (77) and manually inspected. The maximum likelihood phylogenetic analysis was conducted using General Time Reversible (GTR) model with 1,000 bootstraps in MEGA v11 (78) and finalized trees were reconstructed with FigTree v.1.4.3 (79).

### Estimating time to the most recent common ancestor (TMRCA)

For evolutionary dynamic analyses and to determine the age of the most recent common ancestors, the Bayesian Markov chain Monte Carlo (MCMC) method was performed using BEAST v2.6.6 package (80). In these analyses, a GTR + gamma (G) substitution model and a strict clock model were applied (81). MCMC was set to 100,000,000 generations (sampling every 2,500 steps). Log files were analysed in Tracer v1.7.1 to check effective sampling size (ESS) values (>200 indicated sufficient sampling). The maximum clade credibility (MCC) tree was generated in the Tree Annotator v1.8.4, with a default burn-in of 10%. The MCC tree was visualized in the FigTree program v1.4.3 (79).

### Geolocation of USUV strains sequenced in this study

GIS analysis of USUV-positive birds used for sequencing, was performed using ArcGIS ArcMap 10.8.1 (ESRI, Redlands, CA, USA) and open data from GeoBasis-DE/BKG 2022 (82).

### Ethical statement

Veterinarians, wild bird rescue centres, and zoological facilities supplied bird carcasses for necropsy. Residual blood material was available from birds collected primarily for veterinary examination, diagnostic purposes, specific treatments, and the effectiveness of a treatment.

## Results

### USUV genomic sequencing

For WGS, USUV RNA-positive samples were sequenced using ONT (Figs 2 and 3). The median Ct value of the USUV-positive birds was 19.6. S1 Table provides an overview of the total reads and some quality parameters of the sequencing results (coverage, mean read quality, and identity levels) of all samples. The average number of NGS reads obtained from amplicons were approx. 250.000 and genome assembly was performed for all samples with covering >85% of the genome. The total accuracy rates of the USUV genome sequences were 94.2–98.3% with the threshold depth of 100x.

**Fig 2.**
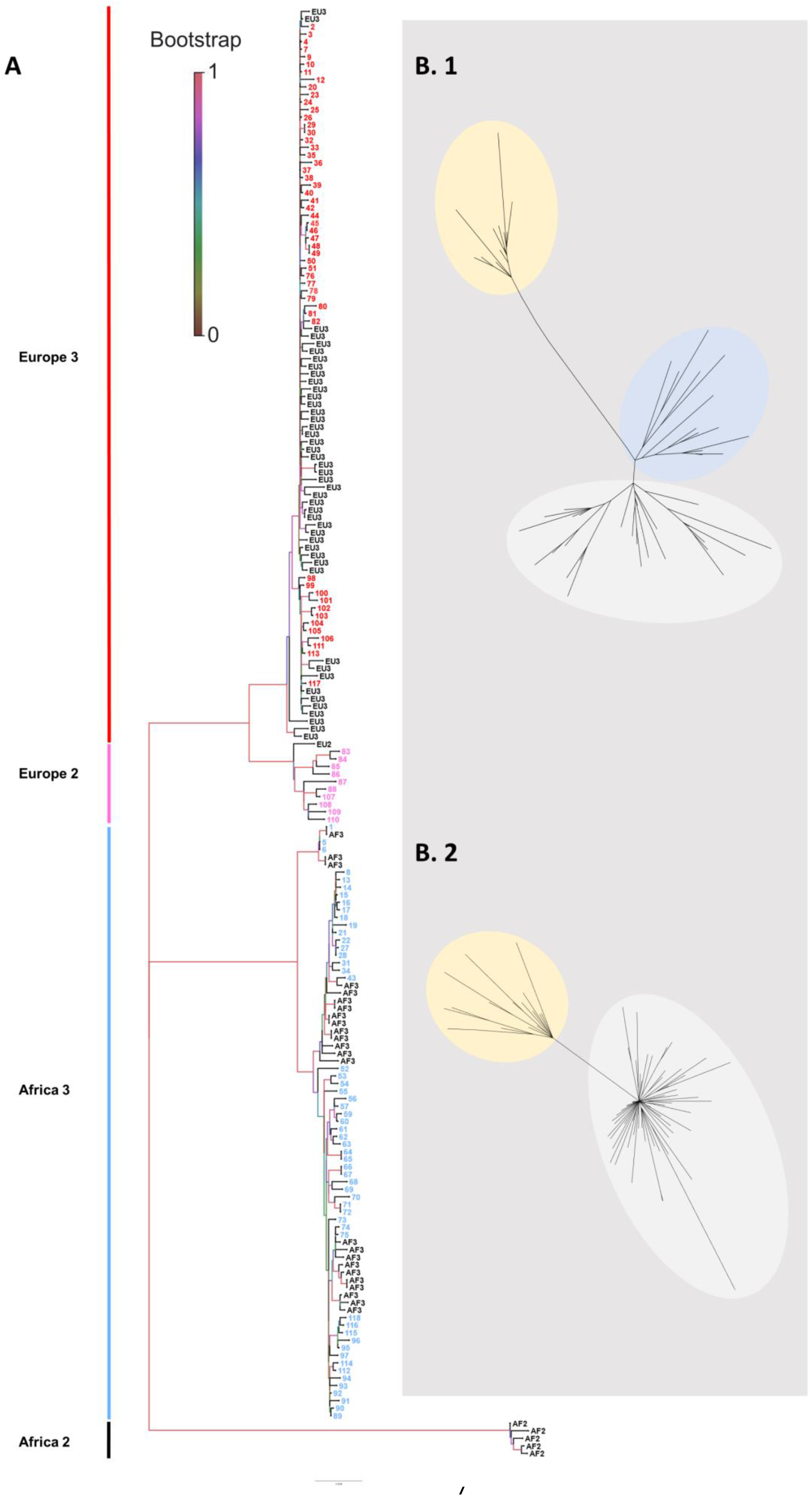
Phylogeny of the sequenced USUV isolates from 2017–2021 (A) Samples were numbered consecutively and coloured according to the lineages. Reference genomes are marked as EU3 (Europe 3), EU2 (Europe 2), A3 (Africa 3), and A2 (Africa 2). Detailed information to each sample number can be found in S1 Table including GenBank accession numbers and the years in which the samples were detected. Scale bars indicate the mean number of nucleotide substitutions per site. (B. 1) Cluster analyses suggesting three subclusters belonging to lineage Europe 3 and (B. 2) two subclusters belonging to lineage Africa 3.

**Fig 3.**
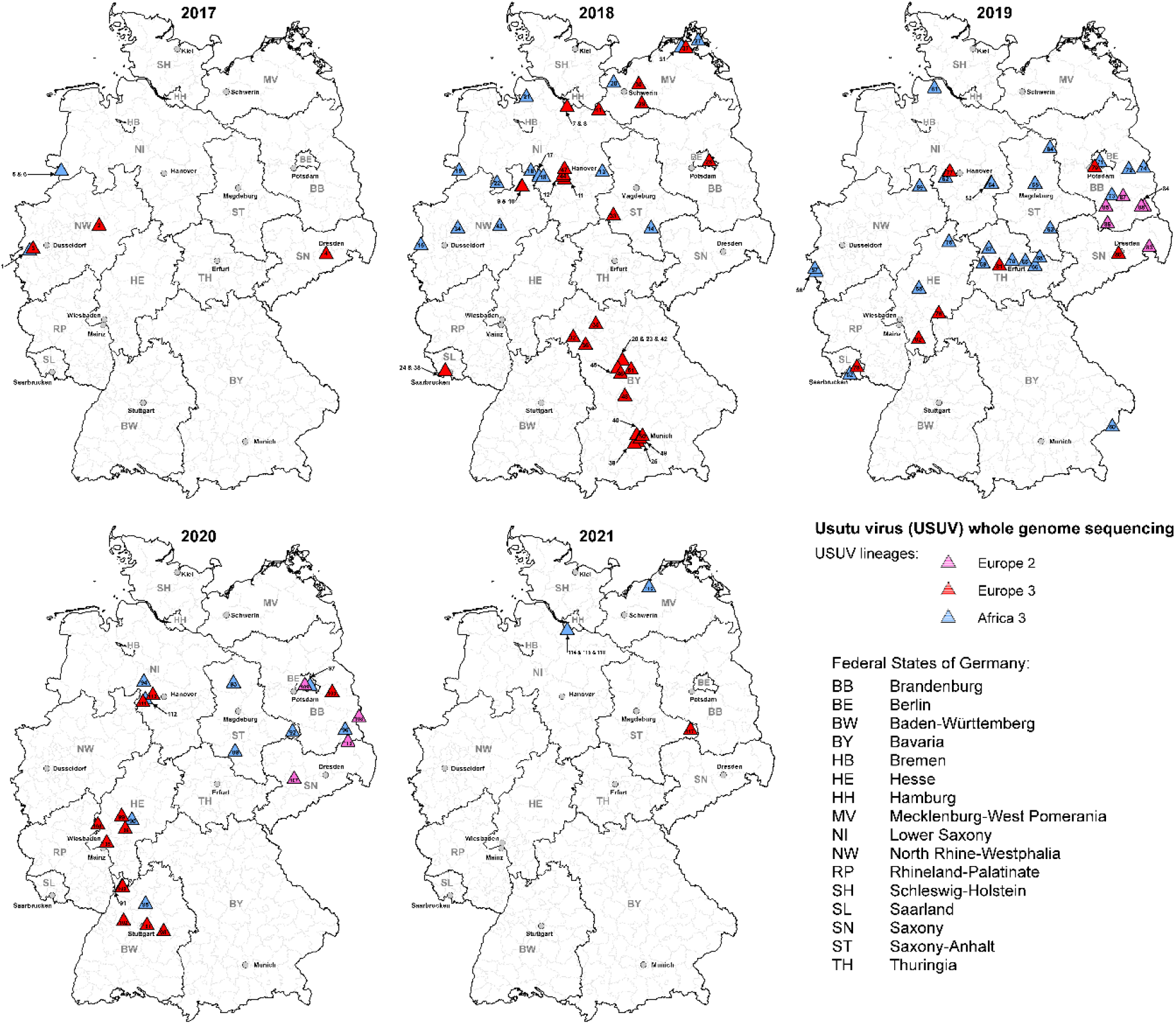
Geographic distribution of whole genome sequences of USUV in Germany from 2017 to 2021. The different USUV lineages are depicted as coloured triangles: pink = Europe 2, red = Europe 3, and light blue = Africa 3 with the appropriate sample number (detailed information to each sample in S1 Table).

### Phylogenetic analysis of USUV in Germany

The phylogenetic analysis displays the co-circulation of USUV lineages Europe 2, Europe 3, and Africa 3 in Germany, with whole genome sequences of lineage Europe 2 only present in 2019 and 2020 (Fig 3). USUV lineage Africa 3 was detected almost as often as Europe 3, while Europe 2 was only sequenced in 9% of the cases (Fig 4). Africa 3 and Europe 3 were distributed throughout the country, whereas Europe 2 was only found in the eastern part of Germany, in the Federal States Berlin, Brandenburg, and Saxony (Fig 3). USUV lineages Europa 5 and Africa 2 were not part of this study.

**Fig 4.**
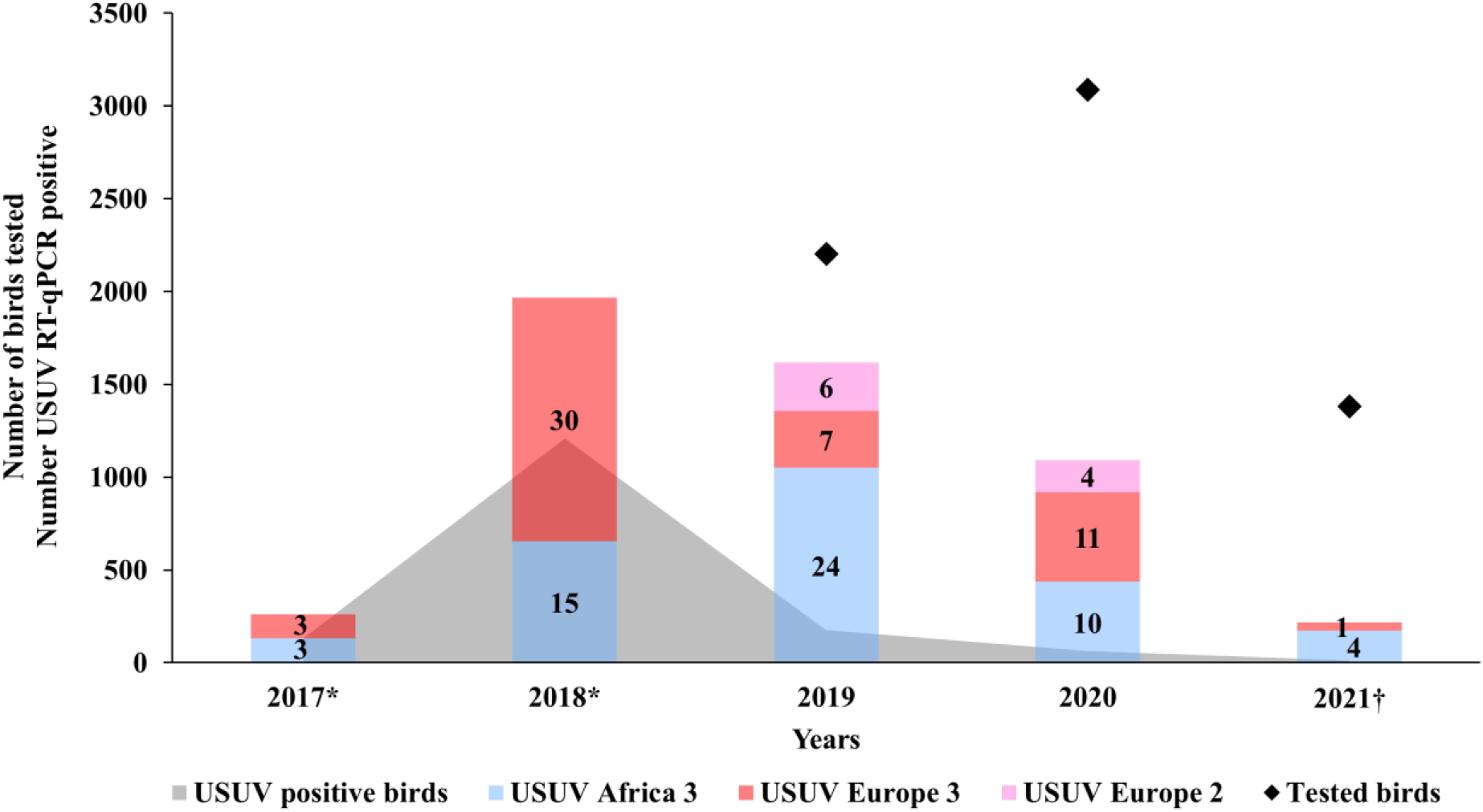
Overview of the distribution of the different USUV lineages in Germany (2017–2021) (depicted in blue, red, and pink with the total number per lineage) with regard to the total number of live and dead birds molecularly tested (depicted with black diamonds) and those tested RT-qPCR positive for USUV (depicted in grey). * Number of tested birds not recorded in 2017 and 2018; ^†^ Results not finalized and only based on dead bird surveillance (updated on 21th February 2023). Bird samples with USUV lineage Europa 5 or Africa 2 were not available for this study.

### Molecular clock phylogeny of USUV lineages detected in Germany

The molecular clock phylogeny of USUV lineages was performed to determine the time to the most recent common ancestors of the USUV lineages Europe 3, Africa 3, and Europe 2. The estimated time to TMRCA of lineage Europe 3 was determined to be around 2008 (between 2006–2010, 95% confidence interval), while the estimated time of lineage Africa 3 was shown to be around 2011 (between 2008–2012, 95% confidence interval), as demonstrated in Fig 5. For Europe 2 TMRCA was calculated to be around 2017 (2015–2019, 95% confidence interval). However, the result for Europe 2 should be considered with caution as the number of available sequences is currently too limited to enable the construction of a fully resolved phylogeny clustering as well as to describe the timing of branching events in phylogenetic trees (S1 Fig).

**Fig 5.**
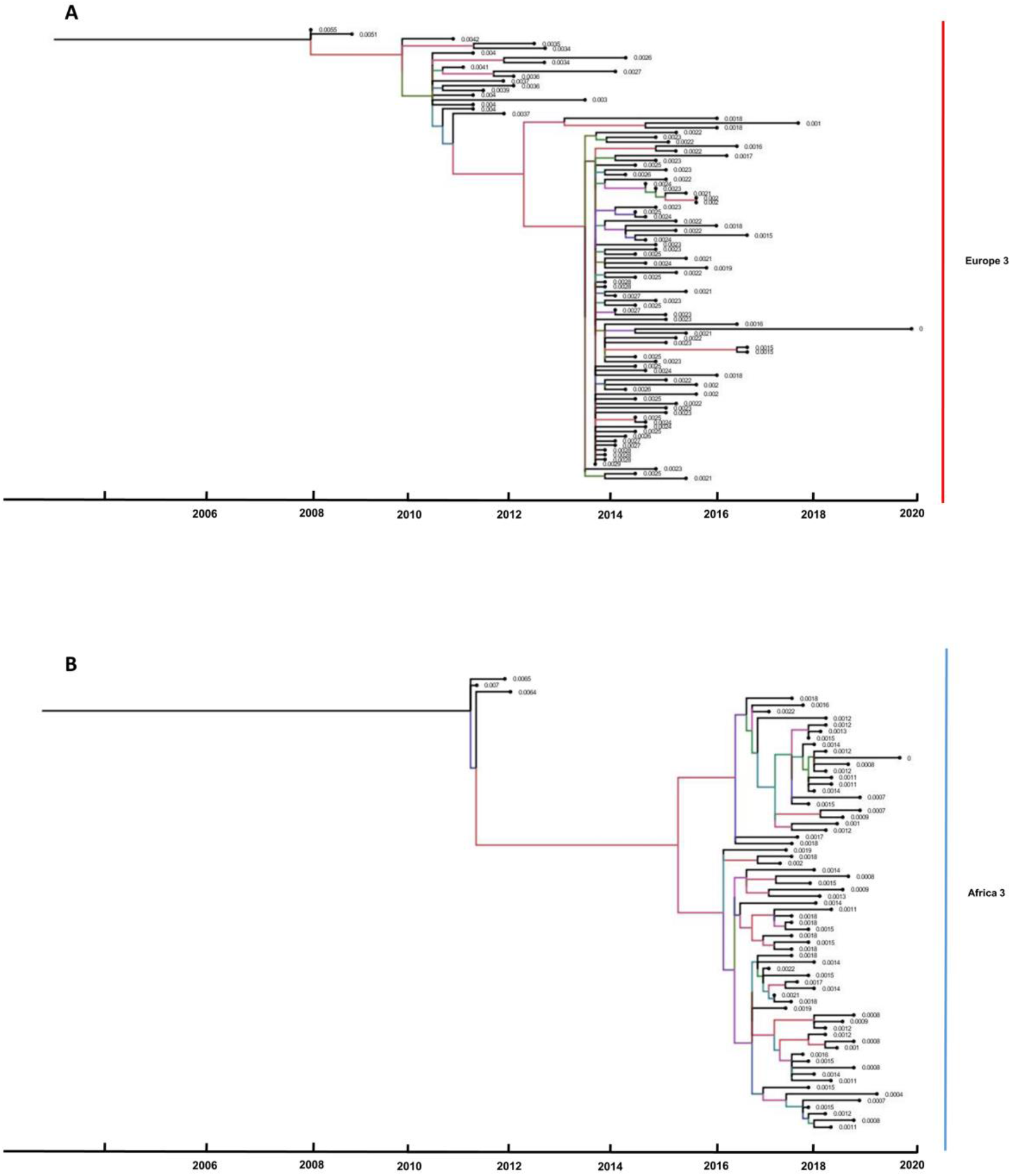
Molecular clock phylogeny of the complete coding sequences of USUV lineages, (A) Europe 3 and (B) Africa 3 detected in Germany. Node bars indicate 95% confidence intervals of the time of TMRCA. The branches are coloured according to the sampling location of their nodes.

## Discussion

The annual reoccurrence of USUV in Germany since 2011 is the cause of ongoing disease and death among wild and captive birds. Regular and stringent genomic surveillance of viral pathogens supports outbreak investigations by providing evidence for their transmission routes and geographic spread. Since only a few whole genome sequences of USUV from Germany are available (67, 47, 8, 83, 33, 15, 49), a better phylogenetic analysis is the next essential step in understanding the spread of USUV in Germany as well as in Europe. A preferred method using third-generation sequencing is a nanopore sequencing approach based on target-enrichment through amplicon generation (84, 67). This recently established protocol (67) was used to gain insight into the distribution and expansion of USUV throughout five consecutive years in Germany. This sequencing technique proved to be sensitive in sequencing the majority of the USUV-positive bird samples and produced good results up to a Ct value of 32.79. The here described study confirms that WGS using the Nanopore platform is suitable in rapidly tracking and detecting ongoing USUV infections in deceased and live birds. Due to the real-time and user-friendly application of the Nanopore sequencing platform, it is a promising tool to supersede partial sequencing in the future. Mass parallelization of sample sets can enable fast-turnaround times without having adverse effects on the platform’s sensitivity in detecting genomic variants (67).

Since the first occurrence of USUV in 2010 in a mosquito pool (one year prior to its detection in birds) (39), five USUV lineages (Europe 2, 3, 5, and Africa 2, 3) (Fig 1) have been described in Germany. Among these, the two USUV lineages Europe 3 and Africa 3 appear to have been the prevailing players in the USUV scenario in the past five years (2017–2021). Therefore, there is an imbalance in the amount of available USUV whole genome sequences for the different USUV lineages. USUV lineage Europe 3 and Africa 3 are predominant compared to only a few whole genome sequences of Europe 2 and Africa 2 and no available whole genome sequences of Europe 5 to date (Ziegler et al. 2016, Cadar et al. 2017). USUV lineage Europe 3, was the first USUV lineage to be detected in Germany, namely in mosquitoes in 2010 (39). However, TMRCA of this lineage is estimated to be about two years prior to its first detection. The 95% confidence interval covers a period from 2006 to 2010 (Fig 5). Similarly, the first detection of USUV lineage Africa 3 occurred in Germany in 2014 (48, 49), yet the TMRCA is estimated to have occurred prior than that, already in 2011 (Fig 5). However, to produce an even more accurate estimate, more data from other geographic areas and earlier years are needed. In contrast, Europe 2 is less frequent in Germany and it was only possible to generate whole genome sequences from 2019 and 2020 (Figs 3 and 4). It should, nonetheless, be noted that partial genome sequences were already generated from samples in 2018, when the lineage was first detected in Germany (45). The TMRCA for Europe 2 was determined in 2017, one year prior to its actual detection. The phylodynamic analyses of TMRCAs therefore provide evidence of a 1-to 3-year lag, respectively between the introduction and the first-case detection of an USUV lineage. By contrast, a large time-lag was described for USUV in the Netherlands, where 7 to 14 years were between the estimated common ancestor and the first detection of USUV lineage Africa 3 and Europe 3, respectively (36). Likewise, the temporal windows determined for the TMRCAs in that publication are broad and can vary depending on the size of the available data set and the set timeframe (36).

The phylogenetic analysis of the USUV whole genome sequences in this study reveal two subclusters for Africa 3. Furthermore, the results of the clock phylogeny (Fig 5 (B)) suggest a long time-lag spanning several years between the first introduction of USUV Africa 3 into the country and its first large-scale occurrence in the avifauna. This could be indicative of silent evolutionary dynamics of the endemic and overwintering lineage Africa 3 (i.e., the first cluster) for several years prior to causing an outbreak with numerous detections (i.e., the second cluster). The occurrence of the outbreak can then be correlated to optimal environmental as well as host and vector conditions driving virus transmission or to the evolution of a more pathogenic strain (i.e., the second cluster). A similar phenomenon was already suggested for WNV by Zehender et al. 2017 (85) and Chaintoutis et al. 2019 (86). They describe that quiet enzootic transmission seasons over several years often proceed virus outbreaks in animals and humans, respectively. Improving the sample matrix (vector and host species) and size as well as the temporal and geographic extent of future surveillance strategies can help on the one hand to detect quiet enzootics and on the other to not miss out on introduction events as well as epizootics. By contrast, the phylogenetic and phylodynamic (Fig 5 (A)) analyses of USUV Europe 3 suggest three subclusters appearing within a shorter time frame. This temporal connection gives the impression of multiple introduction events of USUV Europe 3 into Germany in the past creating the three separate clusters. Alternatively, it is also plausible that the other clusters derived from another undetected USUV Europe 3 isolate endemic in Germany.

In Germany as well as in the Netherlands it, however, remains unclear which influence the absence of USUV surveillance programs had on the first detection of USUV in the two countries. In the case of Germany, the first mosquito monitoring study took place in 2009 at the same sampling site as in 2010, when the first infected mosquito pool was found in Weinheim in the Upper Rhine Valley (39). Prior to 2009, first surveillance efforts were limited to serological investigations, restricted locally (37, 38) and often not primarily focused on flaviviruses (at most testing for WNV (87)). Therefore, it is possible that individual USUV infections occurred before but as they were not accompanied by mass mortality of birds remained undetected. Serological studies prior to 2009 show neutralising antibodies against USUV in wild birds yet no molecular investigations were performed (37, 38). Furthermore, when working with BEAST-analyses one must always keep in mind that a TMRCA is only an estimation of an introduction event based strongly on the quantity and quality of the available data. Caution is always needed when interpreting these results as the inclusion of further samples can reveal different results. This was for example verified by the sequencing of an ancient hepatitis B virus which yielded new data on the evolution of hepatitis B, that was not apparent when only evaluating recent sequences (88).

On a European scale, USUV was isolated for the first time in 2001 in Austria (“Strain Vienna”; USUV Europe 1) (13) and retrospectively in 1996 in Italy (also USUV Europe 1) (14). However, molecular clock analyses revealed that the first entry of USUV (classified as Europe 1) into central Europe was estimated to have occurred already in Spain in the late 20^th^ century, with a virus closely related to USUV from Senegal. Furthermore, the lineages Europe 2 and 3 were calculated to have their origin in Austria in 1993 and Italy in 2007, respectively (15). Since then, USUV has spread throughout Europe with lineage Europe 4 detected in Italy from a sampling set from 2010–2014 with its estimated TMRCA dating back to 2003–2005 (34). Europe 5 was detected for the first time in Germany in 2016 (33) and so far, no TMRCA has been calculated for this lineage. The lineages Africa 2 and 3 descended from multiple independent introductions from sub-Sahara as indicated by analyses from Spain (15).

A few publications have analysed the geographic flow of the USUV genome throughout Europe, with the majority of available whole genome sequences from Italy, the Netherlands, and Germany. Italy is considered an “USUV-donor” to neighbouring countries. Especially north-western Italy appears to have played a key role in the transfer of USUV to central Europe (from Switzerland to Germany to France and Belgium) and eastern Italy to central and eastern Europe (from Austria to Hungary to Serbia) (89). It must however be kept in mind that other publications have described an USUV spread in the other direction, i.e., from Austria to Italy and Germany (15, 34, 4). Analyses performed with USUV sequences from the Netherlands have confirmed an USUV-circulation between the Netherlands, Germany, and Belgium (36). USUV Europe 3 is less frequent in the Netherlands and is most likely periodically re-introduced from neighboring countries. By contrast the prevailing Africa 3 lineage was probably introduced into the Netherlands from Germany in 2016, overwintered there and has since then become enzootic.

Flavivirus transmission dynamics are influenced by environmental and biological factors affecting the host as well as the vector of a virus. The population density of amplifying/reservoir species and the species immune fitness towards specific pathogens also has a significant influence on virus maintenance and spread in the environment (90, 91). Equally, population dynamics of mosquito vectors can play a role in virus transmission and are among others affected by population density, urbanisation, humidity, and temperature (92, 93). The spread of USUV may, therefore, have been favoured by the presence of beneficial environmental conditions for mosquitoes in recent years (33, 94, 95, 8). In 2016, for instance, an exceptionally high activity of USUV infections was observed in birds, correlating with temperature anomalies in September in Western Europe (33). Higher temperatures shorten the extrinsic incubation period (time required for virus replication in the mosquito) which in turn influences the population dynamics of mosquitoes and as a result the vector-host contact rate (92, 96, 90). A similar scenario of optimal weather conditions was observed again in Germany in 2018 with an early humid spring combined with a warm and dry summer (Fig 6) (45, 95). This might have also paved the way for the introduction of WNV into Germany in the same year (97). The extensive USUV outbreaks for example in 2016 and 2018 are in the literature often associated with optimal weather conditions for mosquito-borne virus transmission (98, 90). Especially the fulminant outbreak in 2018 lead to a massive die-off of blackbirds, a species highly susceptible to USUV, and consequently a decline in the species population throughout Germany (45). The observed high seroprevalence of USUV antibodies in 2018 compared to the prior years (45) lets one hypothesize that USUV thereafter faced a higher proportion of non-naïve hosts for its replication cycle. This could help explain the absence of USUV outbreaks of a comparable magnitude in the consecutive years even though the weather conditions continued to encourage arbovirus transmission. This phenomenon was also observed after the USUV outbreak in Austria in 2001, where herd immunity of the wild bird populations protected susceptible species from a severe USUV disease in subsequent years (99).

**Fig 6.**
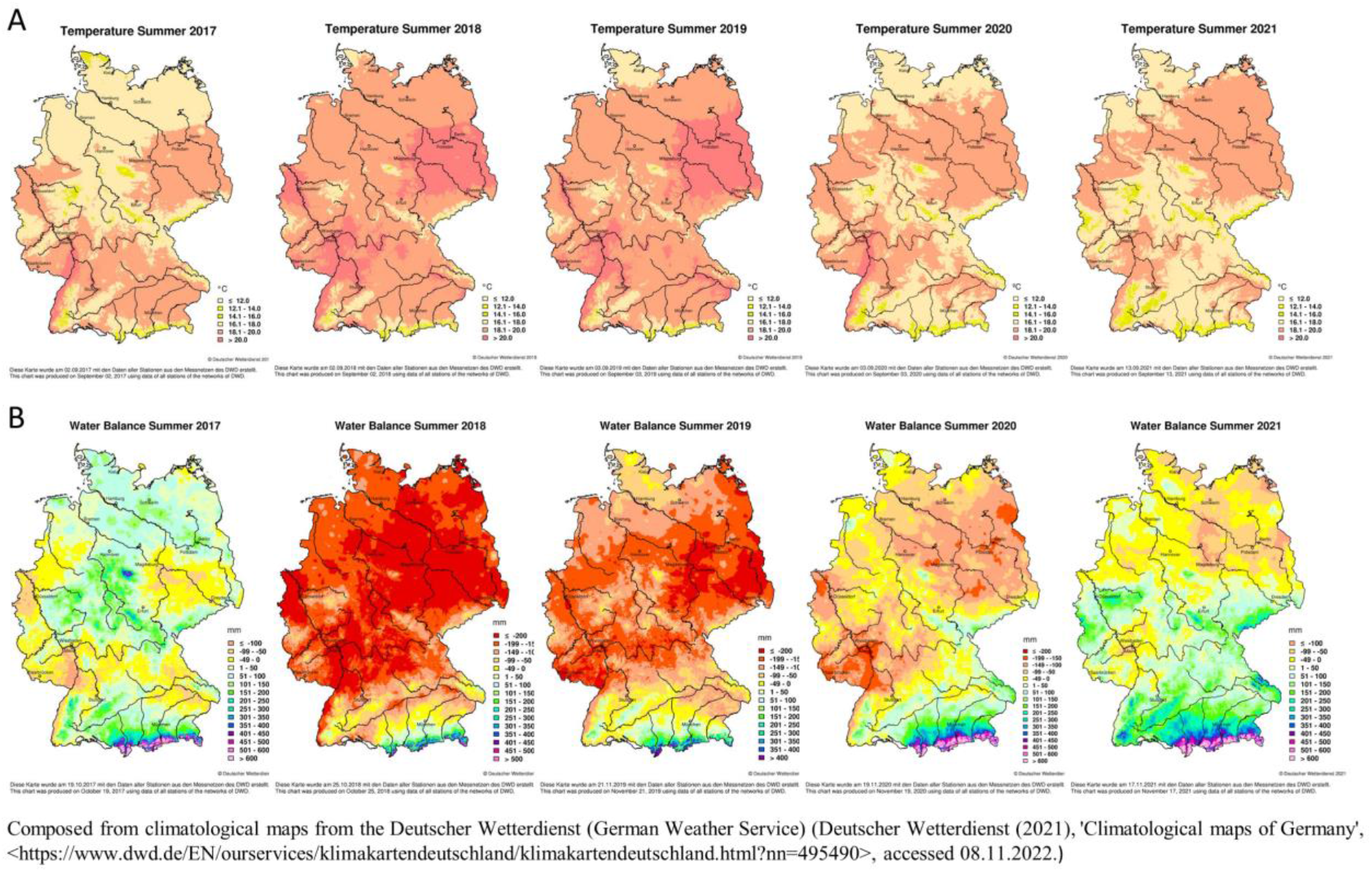
Climatological maps of Germany displaying (A) temperature (in degrees Celsius) and (B) water balance (in millimetre) based on data collected in the summers 2017–2021 (102). Climatological maps were downloaded from the German Weather Service (103).

Arboviruses such as USUV or the closely-related flavivirus WNV can overwinter in a susceptible host and/or vector species. In the past decade, the nationwide German wild bird surveillance network for zoonotic arthropod-borne viruses has consistently provided evidence for the persistence of these arboviruses in the German avifauna (38, 40, 43–45). Similarly, the endemicity of both arboviruses was confirmed in findings from indigenous mosquito species, i.e., *Culex pipiens* (42, 41), that are known to be vector competent for both viruses (92, 100). Increased mosquito breeding, facilitated by high summer temperatures as well as a sufficiently high water balance (Fig 6), can lead to a longer mosquito and virus-transmission season. Taken together the overwintering of mosquitoes infected with flaviviruses (101) and the here discussed evolution of different USUV lineages within the country verify the persistence of the virus in Germany. In addition to optimal weather conditions favouring the spread of endemic USUV lineages in Germany, new introductions from neighbouring countries may have taken place, such as from the Netherlands, Czech Republic or Italy, as well as from long-distance migrants. Newly introduced strains in turn spread with ease under the favourable environmental conditions (Fig 1).

## Conclusion

The study helps to understand the evolution and spread of the major USUV lineages in Germany since their first occurrence in 2010 and to classify the most recent common ancestor for the two most important lineages Europe 3 and Africa 3. For this purpose, a validated protocol was used to efficiently generate whole genome sequences using the Nanopore platform. There was a correlation between the weather conditions and the number of USUV infections detected, with 2018 displaying an exceptionally large USUV epizootic. Using clock phylogenies, the most recent common ancestors were determined for the ubiquitous USUV lineages Europe 3 and Africa 3 in Germany. These results once more emphasize the importance of a stringent surveillance strategy for USUV as well as other flaviviruses as the viruses are to date often first detected two to three years after their calculated introduction.

## Data availability

All data generated or analyzed during this study are included in this published article and its Supporting Information files. Raw sequencing data have been submitted to the NCBI database (XXX) and will be freely accessible if the manuscript is accepted for publication.

## Acknowledgements

The authors are very grateful to Cornelia Steffen and Katja Wittig (both from FLI) for excellent technical assistance. We are also grateful to Patrick Wysocki (FLI) for producing the maps in Fig 3. Furthermore, we thank the staff of the bird clinics/practices, rehabilitation and falconry centres, and ornithologists in the nationwide wild bird surveillance network for zoonotic arthropod-borne viruses who took and sent samples for the annual WNV and USUV monitoring project. Selected samples from the monitoring project were used additionally for whole genome sequencing in this manuscript. Therefore, we thank the following contributors, in particular: Dominik Fischer (Clinic for Birds, Reptiles, Amphibians and Fish, Justus Liebig University Giessen), Sylvia Urbaniak (Raptor Rehabilitation Centre Rhineland), Brigitte Böhm (Tiergesundheitsdienst Bayern e.V.), Mathias Sterneberg (Veterinary practice for raptor medicine, Schüttorf), Karolin Schütte (Wildlife Rescue and Conservation Center, Sachsenhagen), Pierre Grothmann (Wildlife Veterinarian, Geestland), Andreas Kirchhoff (Clinic of Pathology, Gelsenkirchen), Anke Scherer (Nature and Biodiversity Conservation Union (NABU)), Claudia Szentiks (Leibniz Institute for Zoo and Wildlife Research), Niels Mensing (Veterinary practice, Magdeburg), Monika Rinder (Clinic for Birds, Small Mammals, Reptiles and Fish, Centre for Clinical Veterinary Medicine, Ludwig Maximilians University Munich), Kerstin Müller (Department of Veterinary Medicine, Small Animal Clinic, Freie Universität Berlin), Martina Schmoock Wellhausen (Veterinary medicine am Rothenbaum, Hamburg), Heike Weber (Zoological Garden Nordhorn), Florian Hansmann and Anna Schwarz (both from University of Veterinary Medicine Hannover, Department of Pathology), Arne Jung (University of Veterinary Medicine Hannover, Clinic for Birds), Bernhard Böer (Veterinary practice for raptor medicine, Salzgitter), and Heinz-Eike Lange (Veterinary practice, Würselen). We also thank the staff of the wildlife park in Cottbus and Saarbrücken.

## Author contributions

Conceptualization, F.B., C.M.H., B.S., M.H.G., and U.Z.; methodology, F.B., C.M.H., and B.S.; data curation, F.B., C.M.H., U.Z. and B.S.; software, F.B., C.M.H., and B.S.; validation, F.B., C.M.H., B.S., U.Z. and M.H.G.; formal analysis, F.B., C.M.H., U.Z. and B.S.; investigation, F.B., C.M.H., F.M., A.M., S.B., N.S., G.P., S.K., T.S., J.S., C.S., L.H., M.P., and A.H.; resources, F.B., C.M.H., F.M., A.M., S.B., N.S., G.P., S.K., T.S., J.S., C.S., L.H., M.P., A.H., U.Z., and M.H.G.; writing—original draft preparation, F.B., C.M.H., B.S., and U.Z.; writing—review and editing, F.B., C.M.H., B.S., F.M., A.M., S.B., N.S., G.P., S.K., T.S., J.S., C.S., L.H., M.P., A.H., M.H.G., and U.Z.; visualization, F.B., C.M.H, and B.S.; supervision, B.S., U.Z., and M.H.G.; project administration, U.Z. and M.H.G.; funding acquisition, U.Z. and M.H.G. All authors have read and agreed to the published version of the manuscript.

## Funding

This research was funded by the German Federal Ministry of Food and Agriculture (BMEL) through the Federal Office for Agriculture and Food (BLE), grant number 2819113919, by the German Ministry of Education and Research (BMBF) grant number 01KI2026D and by the German Center for Infection Research (DZIF); Project Number TTU 01.804 and 01.808.

## Supporting information

S1 Table. Detailed information on the origin of phylogenetically analyzed USUV from wild and captive birds in 2017 and 2021. Sample numbers are used in Fig 1.

S1 Fig. Molecular clock phylogeny of the complete coding sequences of USUV lineage Europe 2 detected in Germany. Node bars indicate 95% confidence intervals of the time of TMRCA. The branches are colored according to the sampling location of their nodes.

## Notes

### Competing Interest Statement

The authors have declared no competing interest.

